# ODF2 negatively regulates CP110 levels at centrioles/basal bodies to control biogenesis of primary cilia

**DOI:** 10.1101/2023.03.21.533604

**Authors:** Madeline Otto, Sigrid Hoyer-Fender

## Abstract

Primary cilia are essential sensory organelles that develop when an inhibitory cap consisting of CP110 and other proteins is eliminated. Degradation of CP110 by the ubiquitin-dependent proteasome pathway mediated by NEURL4 and HYLS1 removes the inhibitory cap. Here, we investigated the suitability of rapamycin-mediated dimerization for centriolar recruitment and asked whether the induced recruitment of NEURL4 or HYLS1 to the centriole promotes primary cilia development and CP110 degradation. We used rapamycin-mediated dimerization with ODF2 to induce their targeted recruitment to the centriole. We found decreased CP110 levels in transfected cells, but independent of rapamycin-mediated dimerization. By knocking down ODF2, we show that ODF2 controls CP110 levels. Overexpression of ODF2 is not sufficient to promote the formation of primary cilia, but overexpression of NEURL4 or HYLS1 is. Co-expression of ODF2 and HYLS1 resulted in the formation of tube-like structures, indicating an interaction. Thus, ODF2 controls primary cilia formation by negatively regulating the concentration of CP110 levels. Our data suggest that ODF2 most likely acts as a scaffold for the binding of proteins such as NEURL4 or HYLS1 to mediate CP110 degradation.

**Summary:** NEURL4 and HYLS1 mediate the degradation of CP110 to allow cilium formation. We used rapamycin-mediated dimerization with ODF2 to recruit NEURL4 and HYLS1 to the centriole and show that ODF2 controls CP110 levels.

## Introduction

Primary cilia are ubiquitous sensory organelles, essential for the transmission of mechanical and chemical signals toward the cell center, and are, thus, crucial for embryonic and postnatal development, and tissue homeostasis (Badano et al., 2006; Satir and Christensen, 2007; Gerdes et al., 2009; Goetz and Anderson, 2010; Fisch and Dupuis-Williams, 2011; Hoyer-Fender, 2013; Reiter and Leroux, 2017; Higgins et al., 2019). Assembly and disassembly of primary cilia in cycling cells correlate with cell cycle progression and they are predominant in cell cycle arrested, quiescent cells (Wheatley, 1971; Tucker et al., 1979; Quarmby and Parker, 2005; Santos and Reiter, 2008; Plotnikova et al., 2009; Seeley and Nachury, 2010; Kim and Tsiokas, 2011; Avidor-Reiss and Gopalakrishnan, 2013).

The primary cilium is an outgrowth of the basal body that extends from its distal region to eventually project into the cellular environment, while the basal body anchors the cilium inside the cell and to the cell membrane (Ishikawa and Marshall, 2011; Kumar and Reiter, 2021). The basal body itself is a derivative of the older or mother centriole of the pair of centrioles that form the centrosome, which is the main microtubule-organizing center (MTOC) of the cell (Bornens, 2002). Both centrioles of the centrosome, the mother centriole and its descendant, the daughter centriole, differ in age, structure, protein composition, and function. The mother centriole is characterized by the presence of distal (DA) and subdistal (SDA) appendages, and it is the mother centriole that nucleates and anchors microtubules (MTs), and assembles a ciliary axoneme. Initiation of cilium assembly starts with the docking of vesicles at the distal end of the mother centriole, which, additionally, marks the transformation of the mother centriole into the basal body, a process that has to be precisely regulated. In addition to the hundreds of proteins required to build and maintain primary cilia, the onset of cilia formation also requires reorganization and remodelling of the mother centriole (Fliegauf et al., 2007; Pedersen et al., 2008; Ishikawa and Marshall, 2011; vanDam et al., 2019).

At first, after the initiation of centriole duplication, a specific set of proteins is recruited to the nascent daughter centriole. These daughter centriole-enriched proteins (DCPs) include CEP120, Centrobin, and NEURL4 that are also recruited in this order (Zou et al., 2005; Mahjoub et al., 2010; Li et al, 2012). DCPs are removed in the next cell cycle highlighting the conversion of the daughter centriole into a mature centriole and the formation of its characterizing DAs and SDAs. DAs and SDAs are assembled by the sequential recruitment of specific proteins (Yang et al., 2018; Chong et al., 2020). At the tip of DAs locates the protein CEP164, which is involved in the first step in cilia formation, namely the docking of ciliary vesicles (Graser et al., 2007; Schmidt et al., 2012). The assembly of SDAs is centred around ODF2, which is located near the centriole wall (Sullenberger et al., 2020).

Immediately after ciliary vesicle docking, or concurrently with it, the ciliary axoneme is formed by extension from the distal end of the mother centriole/basal body (Sánchez and Dynlacht, 2016). Axoneme extension requires the elimination of the Centriolar Coiled Coil Protein of 110 kDa, in short CP110, and its interacting partner CEP97 from the distal end of the mother centriole (Avidor-Reiss and Gopalakrishnan, 2013; Tsang and Dynlacht, 2013; Nagai et al., 2018). CP110 and its associated partners form a cap at the distal ends of the two centrioles, which prevents microtubule elongation and axoneme extension (Spektor et al., 2007; Schmidt et al., 2009). In contrast to its suppressive role in cilia formation in cultured cells, CP110 is also required for SDA assembly and cilia formation in vivo (Song et al., 2014; Yadav et al., 2016). Its dual role as a suppressor and promoter of ciliogenesis suggests that its optimal cellular level needs to be balanced (Walentek et al., 2016).

CP110 levels are mainly regulated via the ubiquitin-dependent proteasome pathway (D’Angiolella et al., 2010; Li et al., 2013). Ubiquitylation and destabilization of CP110 are promoted by its interactor NEURL4, a protein preferentially located to procentrioles and daughter centrioles (Li et al., 2012). Early during ciliogenesis, in a process that requires mother-daughter centriole proximity, NEURL4 translocates to the mother centriole, which is necessary for the degradation of CP110 and cilia formation (Loukil et al., 2017). NEURL4, which has no ubiquitin ligase activity, is a substrate of the HECT-E3 ligase HERC2, and both are found in a complex with CP110. Thus, the NEURL4-HERC2 complex seems to be responsible for the regulation of CP110 degradation (Al-Hakim et al., 2012). Targeting of NEURL4 to the centrosome was shown to be sufficient to remove CP110 and restore ciliogenesis (Loukil et al., 2017). The removal of the CP110-CEP97 complex then allows the extension of microtubules to form the ciliary axoneme.

Furthermore, additional proteins and CP110 binding partners are involved in the regulation of cilium assembly (Tsang and Dynlacht, 2013). Removal of CP110 and ciliogenesis requires the serine/threonine kinase TTBK2 (Goetz et al., 2012; Cajanek and Nigg, 2014; Oda et al., 2014; Lo et al., 2019). TTBK2 binds to CEP164 at the DAs, which is regulated by the phosphatidylinositol(4)P 5-kinase (PIPKIγ)-mediated depletion of phosphatidylinositol-4-phosphate (PtdIns(4)P) levels at the centrosome/ciliary base (Xu et al., 2016; Wang and Dynlacht, 2018). PIPKIγ, furthermore, interacts and recruits HYLS1 (hydrolethalus syndrome protein 1) to the ciliary base causing activation of PIPKIγ and in turn accelerated depletion of centrosomal PI(4)P and axoneme extension (Dammermann et al., 2009; Chen et al., 2021). In addition to TTBK2, the microtubule (MT)-associated protein/MT affinity regulating kinase 4 (MARK4) is required for the initiation of axoneme extension (Kuhns et al., 2013). MARK4 depletion prevented the exclusion of CP110 from the mother centriole and blocked ciliogenesis.

MARK4 is associated with the basal body and the ciliary axoneme, and interacts with and phosphorylates ODF2 (alias Cenexin), the centriole and sub-distal appendage protein that is mandatory for primary cilia formation and viability (Brohmann et al., 1997). *Odf2*-deficient mice die during preimplantation (Salmon et al., 2006). *Odf2*-deficient centrioles in mouse embryonic carcinoma F9 cells lack appendages and are unable to generate a primary cilium (Ishikawa et al., 2005). Furthermore, a crucial amount of ODF2 at the centrosome/basal body is necessary for the initiation of ciliogenesis (Anderson and Stearns, 2009). ODF2 specifically interacts with the GTPase membrane trafficking regulator Rab8a that facilitates the docking of ciliary vesicles to DAs and the formation of the ciliary membrane (Yoshimura et al., 2007; Wang et al., 2018).

Here, we investigated the suitability of the rapamycin-induced dimerization system for centriolar targeting and asked whether the recruitment of NEURL4 or HYLS1 to centrioles affects cilia formation and CP110 levels. Rapamycin induced dimerization by binding to FKBP12, which contains the FK506 binding protein, and then to the FKBP and rapamycin binding domain (FRB) of mTOR forming a ternary complex (Putyrski and Schultz, 2012). FKBP12 and FRB were fused to either ODF2 or NEURL4 and HYLS1, respectively. We found reduced CP110 levels in transfected cells that were independent of NEURL4 recruitment, indicating that ODF2 is responsible for CP110 removal. We, then, investigated the interrelationship between ODF2 and CP110 and demonstrated that CP110 levels at the centriole and the basal body are controlled by ODF2. Finally, we asked whether primary cilia formation is promoted by overexpression of either ODF2, NEURL4, or HYLS1, or the ODF2-mediated centriolar recruitment of NEURL4 or HYLS1. Our data show that albeit ODF2 controls CP110 levels at centrosomes and basal bodies, primary ciliation was largely not affected. In contrast, we observed a significant increase in ciliated cells by overexpression of either NEURL4 or HYLS1.

Overexpression of both, ODF2 and HYLS1, initiated the formation of intracytoplasmic tubes suggesting the interaction between both proteins that support the formation and/or stabilization of higher-order structures. Our data show that ODF2 is essential for the removal of CP110, and suggest that CP110 levels have to be strictly controlled to allow for cilium formation.

## Results

### ODF2-mediated recruitment of NEURL4 to the centrosome

Removal of the CP110 protein module from the distal end of the mother centriole is essential for axoneme extension and primary cilia formation. The degradation of CP110 is mainly regulated by the ubiquitin-dependent proteasome pathway, in which NEURL4 plays an essential role. We asked whether targeted recruitment of NEURL4 to the distal end of centrioles is sufficient to induce CP110 removal and in turn to promote axoneme extension. We used the rapamycin-system to chemically induce the recruitment of NEURL4 by transfection of plasmids *p138NC(Odf2)::mRFP::FKBP* and *pFRB::ECFP::Neurl4.* Rapamycin binds to both, FKBP and FRB, and thus caused their dimerization. 138NC /ODF2 is intrinsically located in the centrosome and, thus, recruits NEURL4 to the centrosome by rapamycin-induced dimerization. ODF2 was detected by the autofluorescence of the mRFP-tag. Although recruitment of FRB::ECFP::NEURL4 by rapamycin was already visible by the ECFP autofluorescence (Fig. 1A b, f), ECFP-fluorescence was weak. We, therefore, enhanced the fluorescence of FRB::ECFP::NEURL4 by decoration with anti-GFP antibody (Fig. 1B). We found very weak staining of the centrosome for FRB::ECFP::NEURL4 in control cells, which have been incubated with DMSO for 24 hrs (Fig. 1B a-d), but a strong increase at the centrosome by rapamycin-induced dimerization with 138NC(ODF2)::mRFP::FKBP (Fig. 1B). Incubation with 0.2µl of rapamycin for 24 hrs caused centrosomal recruitment of FRB::ECFP::NEURL4 (Fig. 1B e-h) which is much stronger when ten-times as much rapamycin was used but only for 20 min followed by medium exchange and incubation of the cells in standard medium without rapamycin (Fig. 1B i-l).

**Fig. 1.**
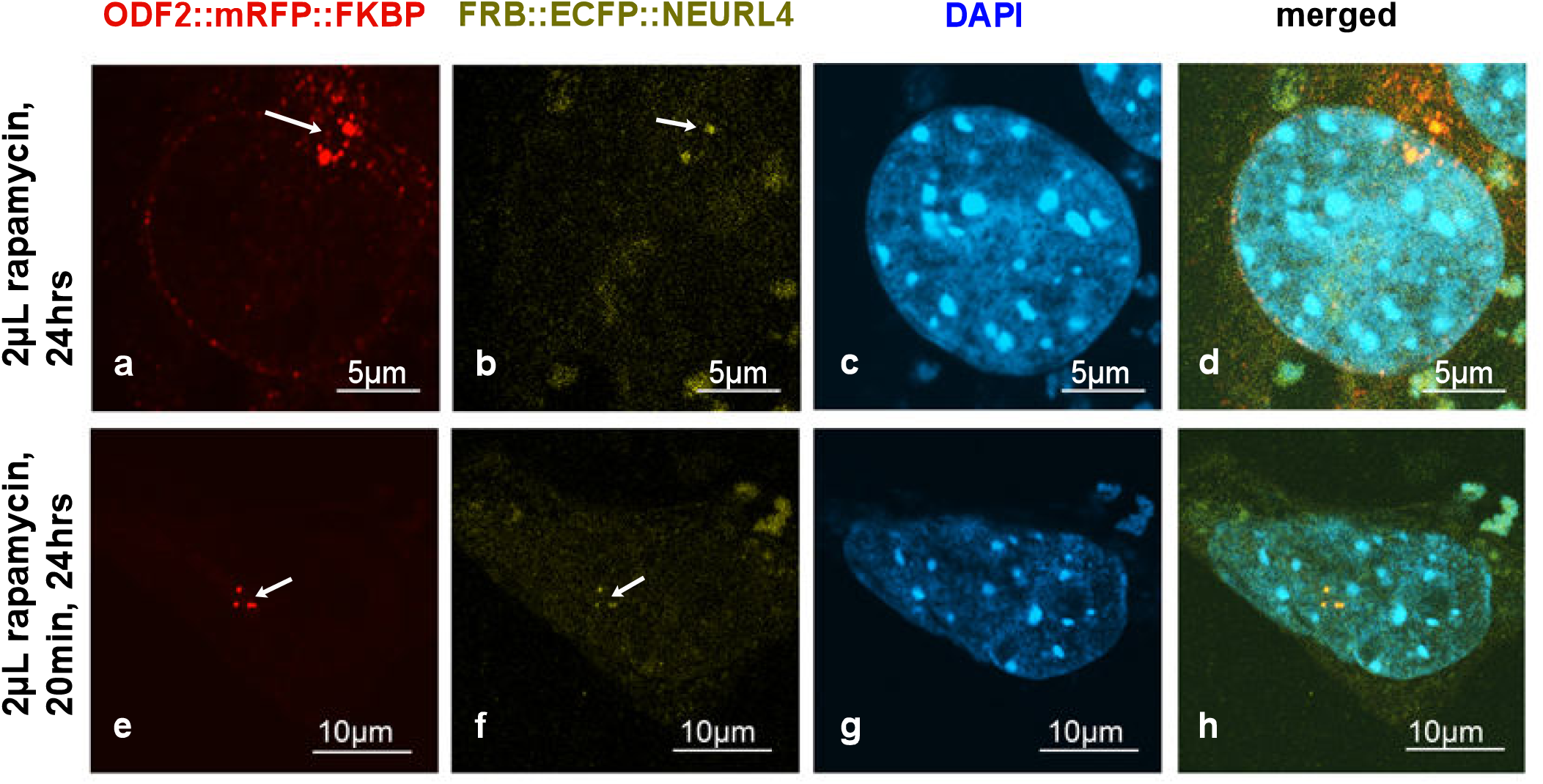
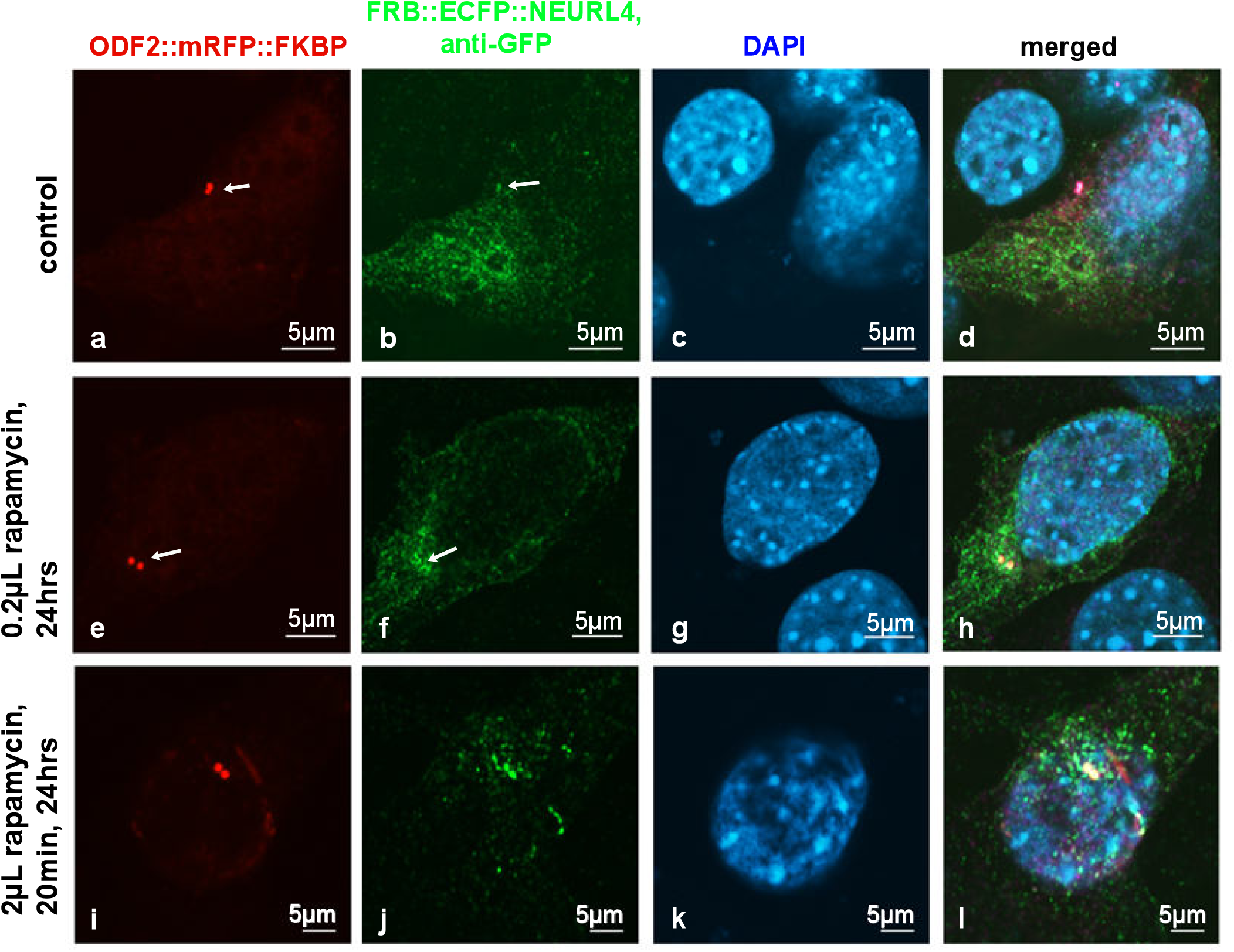
Induced recruitment of NEURL4 to the centrosome by rapamycin-mediated dimerization with ODF2. NIH3T3 cells were transfected with plasmids encoding ODF2 (138NC; 138NC::mRFP::FKBP) and FRB::ECFP::NEURL4. A) Centrosomal recruitment of FRB::ECFP::NEURL4 detected by autofluorescence of either mRFP (a, e, in red) or ECFP (b, f, in green). Nuclear stain with DAPI (c, g, in blue) and merged images (d, h). Scale bares of 5µm (a-d) or 10µm (e-h). Incubation with rapamycin for either 24hrs (2µl of rapamycin, a-d) or 20 min (2µl of rapamycin) followed by medium exchange and incubation without rapamycin for 24 hrs (e-h). B) Decoration of FRB::ECFP::NEURL4 with anti-GFP antibodies. Autofluorescence of the mRFP-tag of ODF2 (a, e, i; in red) at the centrosome (arrows). NEURL4 was immunologically detected with anti-GFP (b, f, j, in green). Nuclear staining with DAPI (c, g, k, in blue) and merged images (d, h, l). Scale bars are 5µm. Incubation with 2µl of DMSO for 24 hrs (a-d) as control. Incubation with rapamycin for either 24hrs (0.2µl of rapamycin, e-h) or 20 min (2µl of rapamycin) followed by medium exchange and incubation without rapamycin for 24 hrs (i-l).

### Rapamycin-induced recruitment of NEURL4 to the centrosome did not enhance the loss of CP110

Since NEURL4 plays an essential role in CP110 removal, we next investigated whether the induced recruitment of NEURL4 to the basal bodies affected the amount of CP110. NIH3T3 cells were transfected with *p138NC/ODF2::mRFP::FKBP* and *pFRB::ECFP::Neurl4* and their dimerization induced by incubation with rapamycin. To identify basal bodies the primary cilium was highlighted by acetylated tubulin staining (Fig. 2A). CP110 was decorated by antibody staining and quantified. Transfected cells without rapamycin incubation and non-transfected cells with or without rapamycin incubation served as controls. We found a significant decrease of CP110 in transfected cells (+T) compared to non-transfected cells (-T) in both cases either without rapamycin-incubation (-R; p=0.001472**) or with rapamycin-induced NEURL4 recruitment (+R; p=0.005132**). No effect on the amount of CP110 was found by rapamycin-incubation of either non-transfected cells (+R-T compared to -R-T; p=0.2759) or transfected cells (+R+T compared to -R+T; p=0.968234) (Fig. 2B). Our data, thus, show that ODF2-mediated, rapamycin-induced recruitment of NEURL4 to the basal body did not cause a decrease in CP110. Furthermore, the data indicate that ODF2 is responsible for the loss of CP110.

**Fig. 2.**
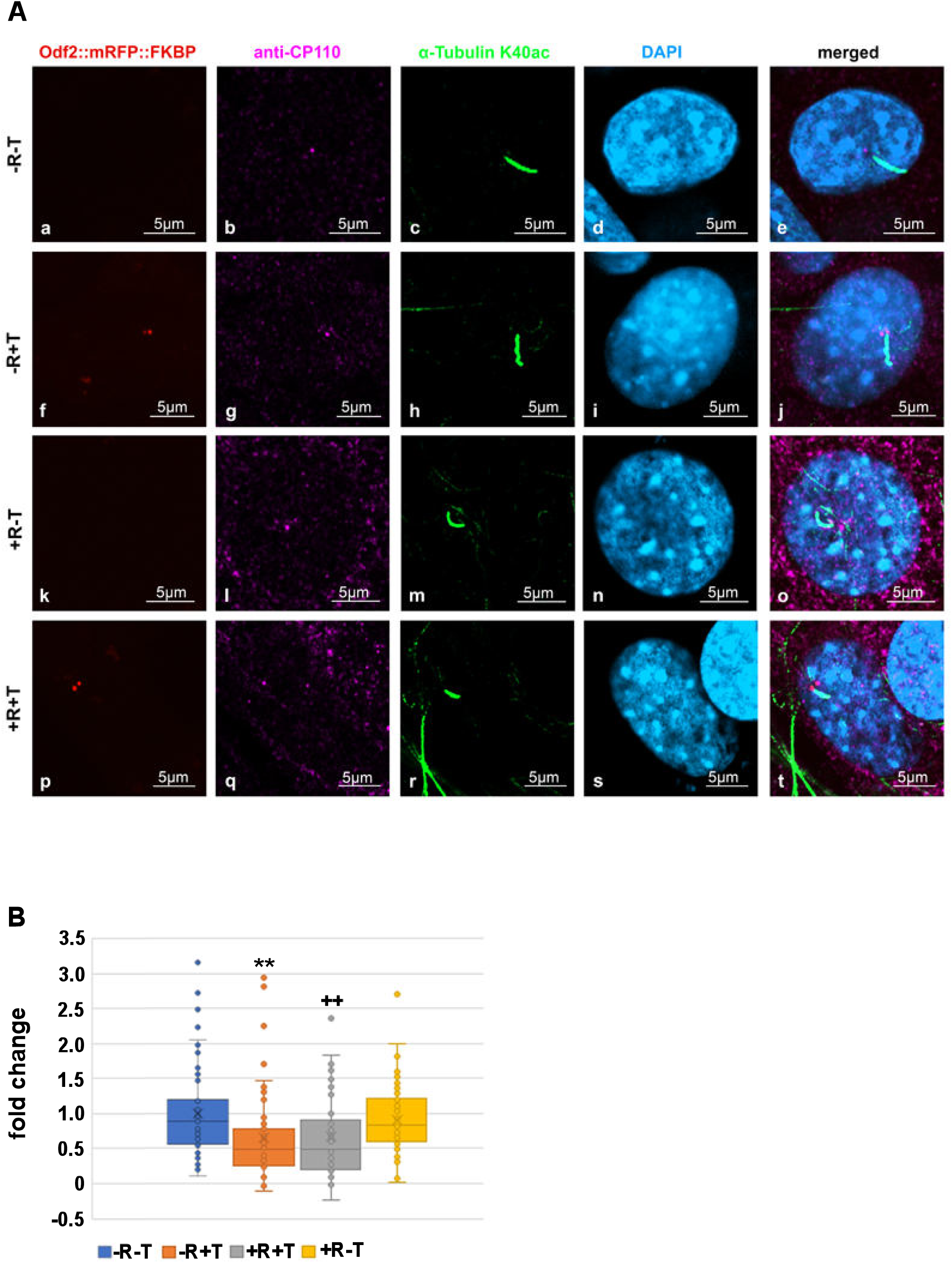
Increased amount ODF2 at the basal body but not rapamycin-induced recruitment of NEURL4 caused loss of CP110. NIH3T3 cells were transfected with *p138NC/ODF2::mRFP::FKBP* and *pFRB::ECFP:::Neurl4* (+T) and incubated either with (+R) or without rapamycin (-R). Non-transfected cells (-T) served as controls. A) Detection of the basal body in transfected cells by autofluorescence of the mRFP-tagged ODF2 (a, f, k, p; in red). Anti-CP110 staining in b, g, l, q (in magenta). Basal bodies were identified by their association with the ciliary axoneme decorated for acetylated tubulin (c, h, m, r; in green). DAPI staining of nuclei (d, i, n, s; in blue) and merged images (e, j, o, t). Scale bars are 5µm. B) CP110 was identified by antibody staining and the CP110 area was captured by confocal imaging. CP110 intensities of individual basal bodies were quantified, followed by background subtraction, which was calculated as the average intensity of four different neighbouring areas, and the corrected CP110 intensity was divided through the area. The average of the calculated CP110 intensities of the basal bodies of control cells (non-transfected and not incubated with rapamycin, -R-T) served as the reference to which all calculated CP110 intensities were related, giving the fold changes. In transfected cells, the average CP110 intensities were 0.658x (-R+T) and 0.6629x (+R+T) compared to the control (-R-T, 1x), whereas rapamycin-incubation in non-transfected cells (+R-T) had no effect showing an average intensity of 0.9124x. A significant decrease of CP110 was found in transfected cells in the absence of rapamycin (-R+T compared to -R-T; p=0.001472**) as well as in the presence of rapamycin (+R+T compared to +R-T; p=0.005132**^++^**). (Student’s T-test two-tailed, homoscedastic). N individual quantifications were performed: n=92 (-R-T), n=54 (-R+T), n=88 (+R-T), n=50 (+R+T). p<0.01** and **^++^**.

### Distribution of ODF2 and CP110 at the centrosome and the basal body

Our data indicated that the removal of CP110 is controlled by ODF2. We, thus, looked first at the distribution of both, ODF2 and CP110, in centrosomes and basal bodies in NIH3T3 cells. Decoration of the cilium by anti-acetylated tubulin staining served to discriminate between centrosomes and basal bodies. To allow for concurrent detection of ODF2 and CP110, cells were transfected with a plasmid encoding ODF2 fused to mRFP (*p138NC::mRFP::FKBP*) for autofluorescence-detection of ODF2 and immunologically stained for CP110 and acetylated tubulin. ODF2 predominantly labelled the mature centriole in the centrosome and the basal body, which is located at the base of the primary cilium, whereas the associated daughter centriole, although decorated with ODF2, showed a much weaker staining (Fig. 3 a, f). CP110 essentially decorates only one of the two centrioles (Fig. 3 b, g). Thus, an inverse relationship of the amount of both proteins seems to exist in that a large amount of ODF2 is associated with low CP110 levels and vice versa corroborating our previous data.

**Fig. 3.**
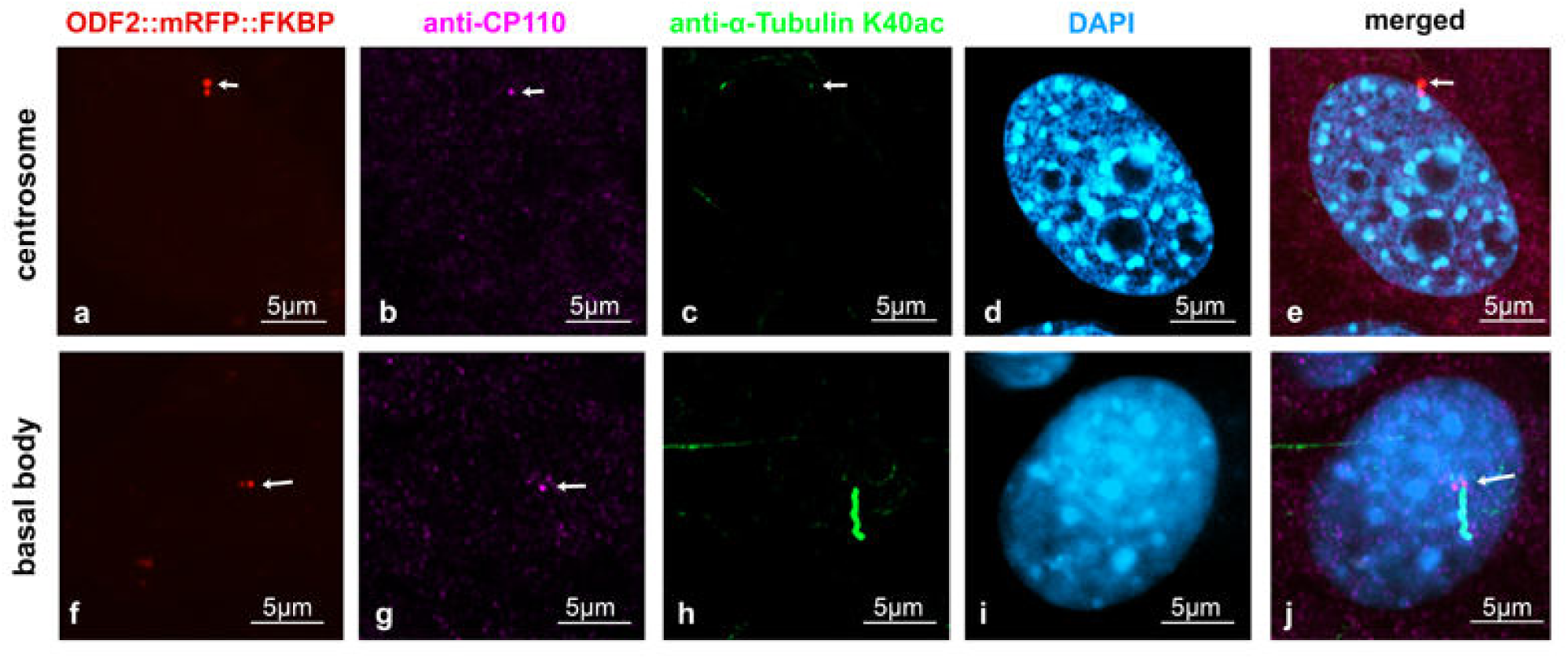
ODF2 and CP110 levels are inversely correlated at centrosomes and basal bodies. NIH3T3 cells were forced to express ODF2 fused to mRFP (138NC/ODF2::mRFP::FKBP) (a, f, red), which was detected by the autofluorescence of mRFP, whereas CP110 (anti-CP110, b, g, pink) and acetylated tubulin (anti-α-tubulin K40ac, c, h, green) were detected by antibody staining. The centrosome was decorated for acetylated tubulin (c, arrow), and acetylated tubulin decorated the primary cilium at the basal body (h). Nuclei were stained with DAPI (d, i, blue). Merged images in e, j. Scale bars are 5µm. Arrows highlight centrosomes and basal bodies.

### ODF2 promotes depletion of centrosomal CP110

To investigate whether the amount of CP110 is affected by ODF2, we modified the amount of ODF2 by either depletion or overexpression and quantified CP110 at the centrosome. Cells were transfected with *pCentrin::Cherry* for decoration and identification of the centrosome, and ODF2 depleted by co-transfection of either a short hairpin plasmid targeting *Odf2* (sh3) or *Odf2* siRNA. For rescue, *hCenexin* was co-transfected. As controls, either the non-targeting control plasmid (K07) or a scrambled control siRNA were transfected. Successful depletion or rescue of ODF2 was verified by quantification of ODF2 at the mother centriole (Fig. 4A). We found a reduction of ODF2 by *sh3*-mediated depletion to ∼0.03-fold compared to the control plasmid *K07*, and an increase by co-transfection of *sh3*+*hCenexin* to∼0.6-fold of the control. Depletion of ODF2 by *Odf2* siRNA caused a reduction to ∼0.03-fold compared to the scrambled control siRNA and an increase to ∼0.53-fold compared to the scrambled control siRNA in the rescue experiment. Thus, ODF2 was significantly depleted by both, either the short hairpin plasmid *sh3* (p=3.73×10^-10^ ****) or by *Odf2* siRNA (p=3.48×10^-10^ ****) compared to the respective controls, either *K07* or scrambled siRNA. Additionally, co-transfection of *hCenexin*-plasmid with either the *sh3* plasmid or the *Odf2* siRNA rescued the respective depletion (*sh3*+*hCenexin* to *sh3*: p=8.135×10^-14^ ****; *Odf2* siRNA+*hCenexin* to *Odf2* siRNA p=7.37×10^-9^ ****). Significance was calculated by Student’s T-test (two-tailed, homoscedastic).

**Fig.4.**
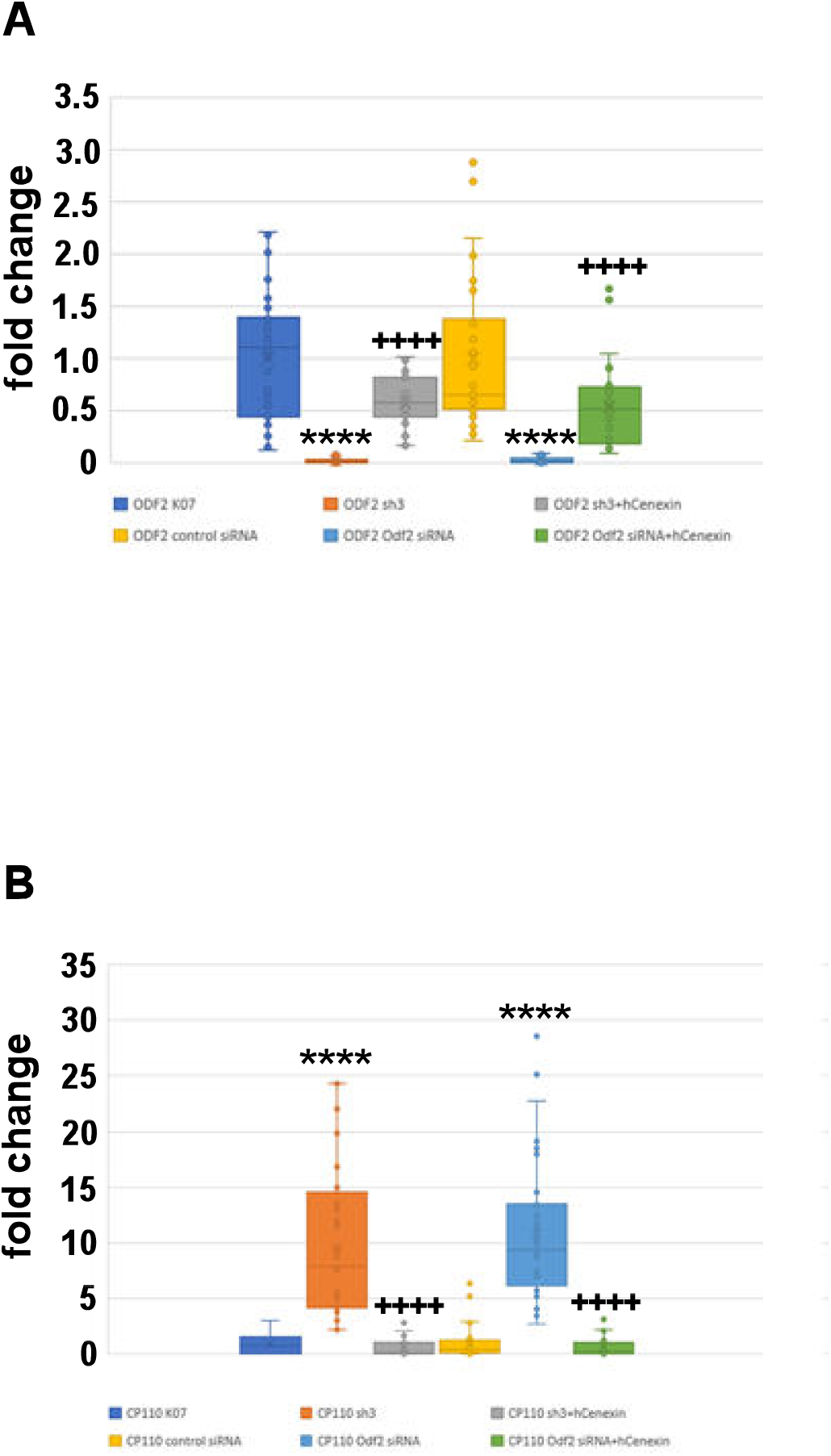
Inverse correlation between the amount of ODF2 and CP110 at the centrosome. ODF2 was depleted by transfection of either the short hairpin plasmid *sh3* or siRNA. Depletion was rescued by co-transfection of *hCenexin* (*sh3*+*hCenexin* or *Odf2* siRNA+*hCenexin*). *K07* or control siRNA served as the controls. A) Centrosomal quantification of ODF2 revealed significant ODF2-depletion by either *sh3* or *Odf2* siRNA transfection, and a successful rescue by co-transfection of *hCenexin*. Three biological replicates were used for analyses with n individual centrosomes: *K07* n=48, *sh3* n=19, *sh3*+*hCenexin* n=35, control siRNA n=35, *Odf2* siRNA n=29, *Odf2* siRNA+*hCenexin* n=31. P<0.0001 **** to the respective control, either K07 or control siRNA, p<0.0001^++++^ to the respective knockdown, either *sh3* or *Odf2* siRNA. B) Quantification of CP110 at the centrosome in ODF2-depleted cells. Three biological replicates were used for analyses with n individual measurements: *K07* n=17, *sh3* n=44, *sh3*+*hCenexin* n=22, control siRNA n=33, *Odf2* siRNA n=40, *Odf2* siRNA+*hCenexin* n=28. P<0.0001 **** to the respective control, either K07 or control siRNA, p<0.0001^++++^ to the respective knockdown, either *sh3* or *Odf2* siRNA.

We then quantified CP110 at the centrosome in cells either depleted or rescued for ODF2 (Fig. 4B). As before, the centrosome was identified by Centrin::Cherry fluorescence. We found an increase in centrosomal CP110 of ∼10x in ODF2-depleted centrosomes by either *sh3* (p=8.789×10^-7^****) or *Odf2* siRNA transfection (p=2.15×10^-13^****) compared to the respective controls, either *K07* or control siRNA transfection. The rescue experiments caused an ∼0.6x amount of CP110 in *sh3*+*hCenexin* transfected centrosomes (p=1.1787×10^-8^****) and an ∼0.7-fold increase in *Odf2* siRNA+*hCenexin* transfected cells (p=1.49×10^-12^****) when compared to the respective controls either *K07* or control siRNA (Student’s T-test two-tailed, homoscedastic). Our data show that ODF2 depletion correlated with an increase of CP110 at the centrosome.

Exemplary pictures used for the quantification of ODF2 and CP110 by ODF2 knockdown are shown in Fig. 5 and 6. All pictures were captured by identical settings and no further processing was done to allow for both a visual estimation as well as a correct protein quantification. Centrosomes were identified by Centrin-Cherry auto-fluorescence. Protein quantification of Centrin-Cherry-labelled centrosomes only also ensured that only transfected cells were included in the analyses. A marked decrease in ODF2 at the centrosome is observed by *sh3*-or siRNA-mediated knockdowns (Fig. 5 b, f, and n, r), as is the rescue of ODF2 by con-transfection of *hCenexin* (Fig. 5 f, j, and r, v). The ODF2 knockdown correlated with increased CP110 at the centrosome (Fig. 6 b, f, and n, r).

**Fig. 5.**
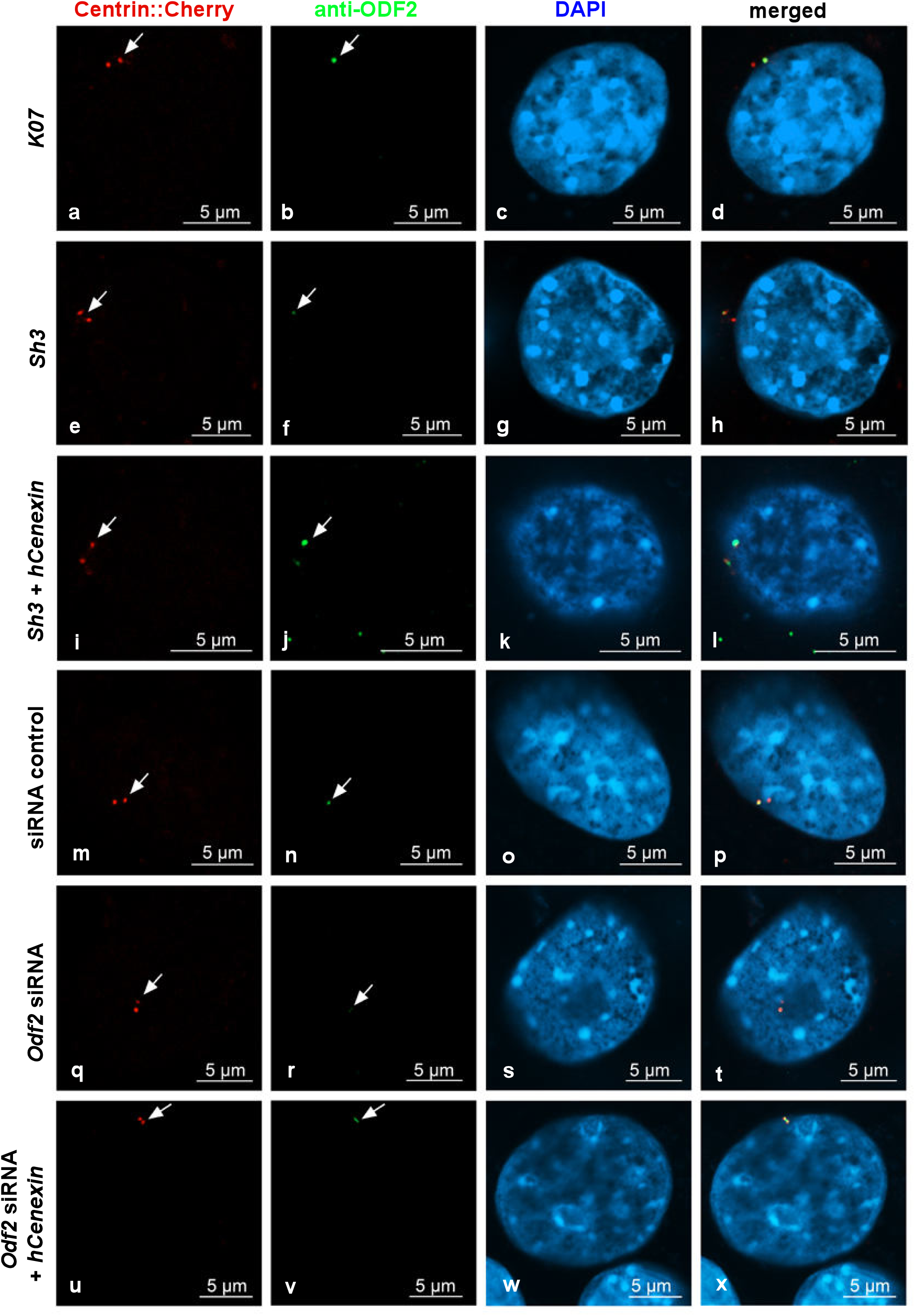
Decrease of centrosomal ODF2 by both, either *sh3*-or *Odf2* siRNA-mediated knockdown. Transfected cells were identified by the autofluorescence of Centrin-Cherry-tagged centrosomes (a, e, i, m, q, u, in red). ODF2 was detected by antibody staining (b, f, j, n, r, v, in green). Nuclear stain with DAPI (c, g, k, o, s, w, in blue). Merged images in d, h, l, p, t, x. Cells were transfected with either the control plasmid *K07*, the short hairpin plasmid *sh3,* or both, *sh3*+*hCenexin,* for rescue. Additionally, the knockdown of ODF2 was also achieved by transfection of *Odf2* siRNA and rescued by co-transfection of *Odf2* siRNA and *hCenexin*. Control siRNA transfection served as the control experiment. Scale bars are 5µm. Arrows highlight the centrosomes.

**Fig. 6.**
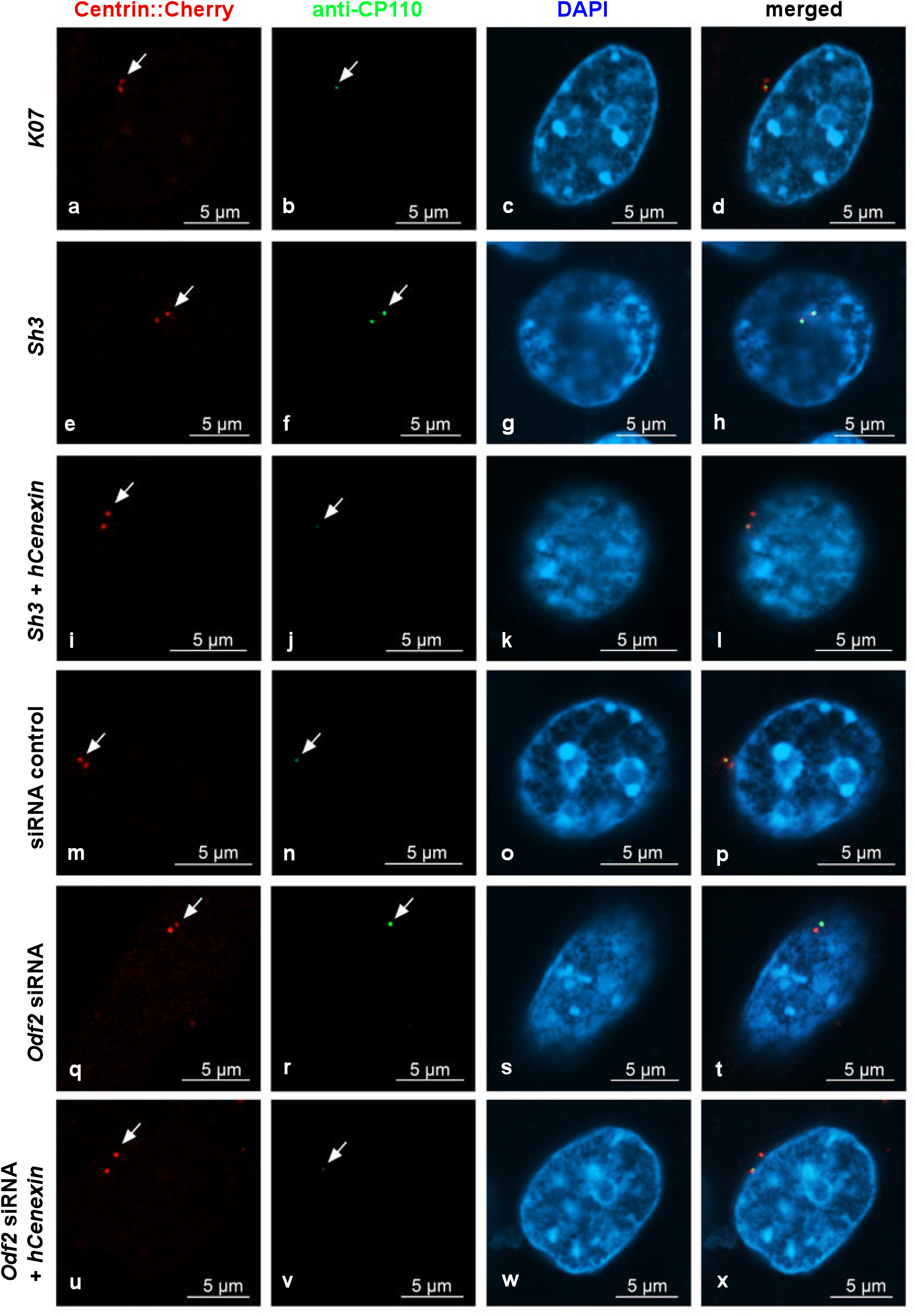
ODF2 knockdown correlated with increased CP110 at the centrosome. Detection of transfected cells by the autofluorescence of Centrin-Cherry-tagged centrosomes (a, e, i, m, q, u, in red). CP110 was detected by antibody staining (b, f, j, n, r, v, in green). Nuclear stain with DAPI (c, g, k, o, s, w, in blue). Merged images in d, h, l, p, t, x. For ODF2 knockdown, cells were transfected with either the short hairpin plasmid *sh3,* and for rescue with both, *sh3*+*hCenexin*. Additionally, the knockdown of ODF2 was also achieved by transfection of *Odf2* siRNA and rescued by co-transfection of *Odf2* siRNA+*hCenexin*. Transfection of the control plasmid *K07* or control siRNA served as the reference experiments. Scale bars are 5µm. Arrows highlight the centrosomes.

### Enrichment of ODF2 at the basal body

ODF2/Cenexin is a marker protein for the mature centriole and the basal body and is mandatory for primary cilia formation. Initiation of cilia formation requires the transformation of the mature centriole into a basal body, a process including structural and compositional remodelling which might also encompass ODF2. We, therefore, asked whether the formation of cilia correlated with an increase in ODF2 and, thus, quantified the amount of ODF2 in centrosomes and basal bodies in both, cycling cells and serum-starved cells that have been cultivated in either standard medium (NM) or serum-depleted medium (SSM), respectively. Discrimination between centrosomes and basal bodies was achieved by the decoration of the axoneme of the primary cilium for tubulin acetylation (Fig. 7A, in green). Centrosomes were identified by the detection of ODF2, located in both centrioles and represented as twin dots, but without an attached axoneme (Fig. 7A, in red). We found a significantly increased ODF2 concentration in basal bodies compared to centrosomes in both conditions (Fig. 7B). In proliferating cells, cultivated in standard medium with 10% FCS, ODF2 is ∼1.5-fold enriched in basal bodies compared to centrosomes (p=5.662×10^-5^ ****), whereas in serum-starved cells enrichment of ODF2 in basal bodies is ∼1.2-fold compared to centrosomes (p=0.04186 *). Enrichment of ODF2, thus, correlated with the transformation of the centrosome into the basal body and cilium formation.

**Fig. 7.**
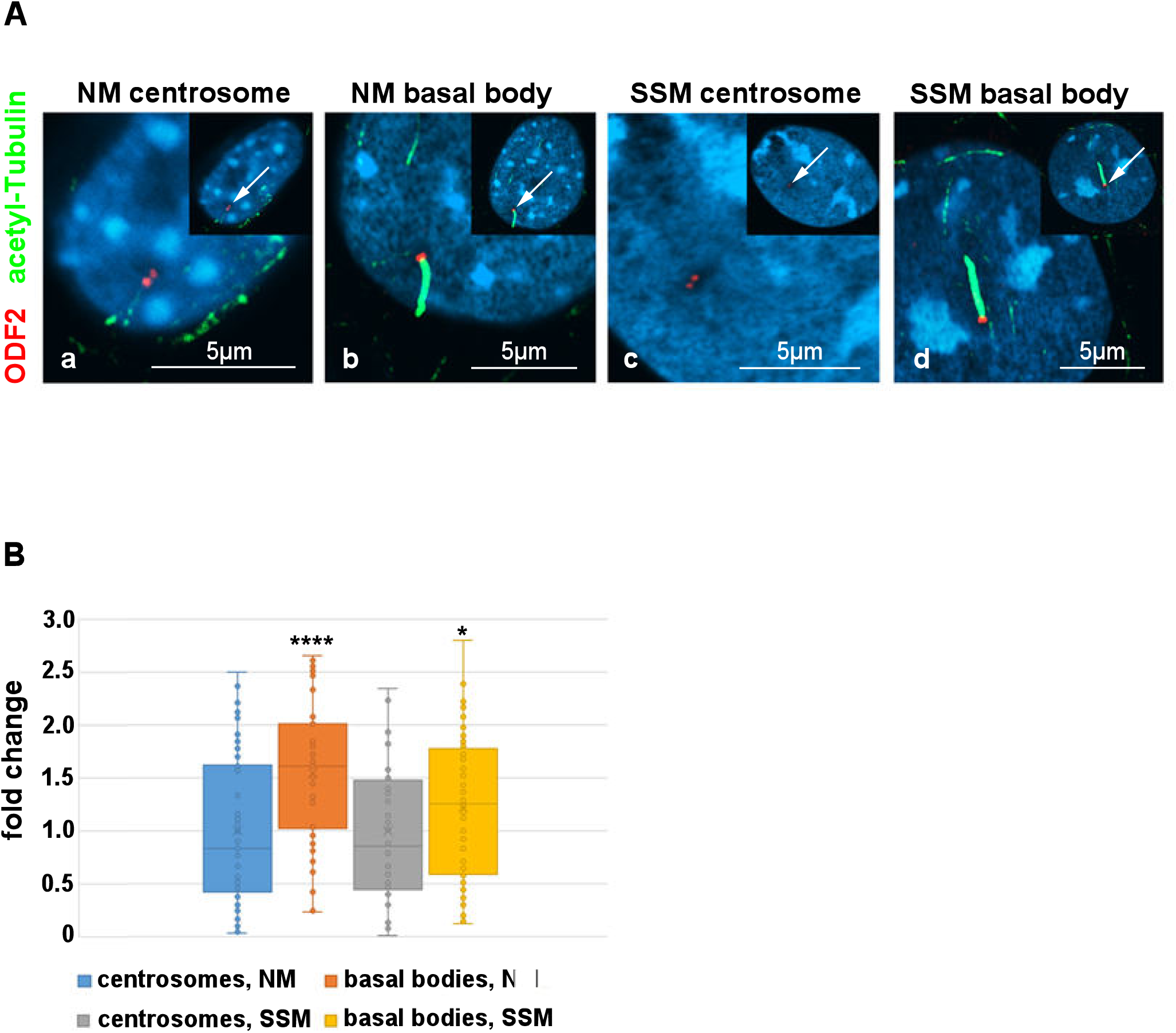
ODF2 is enriched in basal bodies. A) Detection of endogenous ODF2 (in red) in centrosomes and basal bodies of NIH3T3 cells. Cells were cultivated in either normal medium (NM) or serum starvation medium (SSM) and centrosomes (a, c) and basal bodies (b, d) were detected by anti-ODF2 (in red) and anti-acetylated tubulin (in green) decoration. Basal bodies were identified by the presence of the primary cilium, which was decorated by anti-acetylated tubulin staining (in green). Nuclear staining with DAPI. All merged images. The insets show an overview of the whole cell with the enlarged area highlighted by an arrow. Bars are 5µm. B) ODF2 was enriched in basal bodies as compared to centrosomes under both cultivation conditions either normal medium or serum starvation medium. Quantification was done by Fiji. For each quantification, the average intensity of the centrosomal/basal body background (using always four different areas) was subtracted from the intensity of ODF2 staining and the corrected intensity related to the measured area thus giving the relative quantity of ODF2. For the fold change calculation, the relative quantities of ODF2 were related to the average relative quantity in centrosomes either in NM or SSM, respectively. Three biological replicates with n measurements: centrosome in NM n=52, basal body in NM: n=48, centrosome in SSM n=44, basal body in SSM n=66.

### Overexpression of ODF2 did not promote cilia formation

Our data showed that ODF2 levels are increased in basal bodies compared to centrosomes and that ODF2 levels correlated inversely with CP110 levels. Thus, ODF2 regulated the elimination of the inhibitory cap at the distal end of the mother centriole to allow cilia to grow out. We, therefore, asked whether increased expression of ODF2 would stimulate primary cilia formation. Cells were cultured in either standard medium or serum-deprived medium and the percentage of ciliated cells was determined by decorating for ARL13b. We found an increase from ∼30% ciliated cells in standard medium to >70% in serum-deprived medium. However, no significant differences, but rather slight decreases, were observed between control cells and cells transfected with either *hCenexinΔGFP*, *hCenexin::GFP* or *p138NC::mRFP::FKBP,* whether incubated with or without rapamycin (Table 1). We concluded that neither an increase in ODF2 nor incubation with rapamycin is sufficient for primary cilia formation.

**Table 1.**
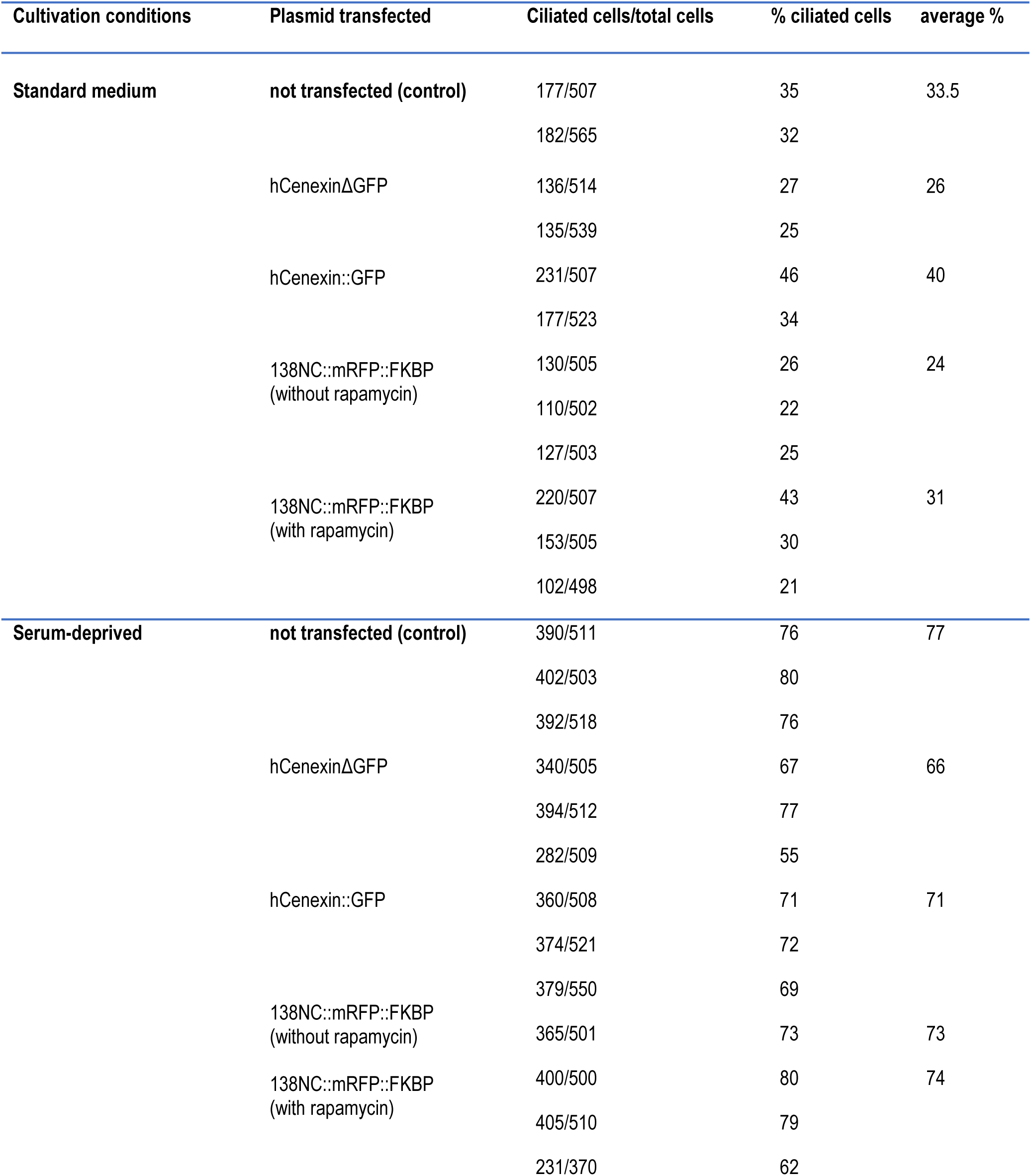
Overexpression of ODF2 is not sufficient for primary cilia formation. Cells were transfected with the indicated plasmids, cultivated in either standard medium or serum-deprived medium, and *p138NC::mRFP::FKBP*-transfected cells were treated either with or without rapamycin. Primary cilia were decorated for ARL13b, manually counted by focusing through all focal planes, and the percentage of ciliated cells was calculated.

### Ciliogenesis and Neurl4 expression

Next, we asked whether the rapamycin-induced dimerization with ODF2 causing the recruitment of NEURL4 to the centriole, promoted cilia formation. Cells were transfected with *p138NC::mRFP::FKBP* and *FRB::ECFP::Neurl4* and either incubated without or with rapamycin followed by immunodecoration of ARL13b. An average of 12% ciliated cells were found in the untreated control cells (ciliated cells/total cells counted: 54/506, 78/516, 61/519, 57/508, four biological replicates). When transfected with both plasmids around 10% of transfected cells are ciliated without rapamycin-incubation (transfected cells with cilium/total transfected cells counted: 2/102, 26/104, 3/104, three biological replicates), and rapamycin-induced dimerization caused an increase to 13% (transfected cells with a cilium/total transfected cells counted: 25/110, 14/103, 3/103, three biological replicates). For cilia counting, only transfected cells that were identified by the autofluorescence of the 138NC::mRFP::FKBP-protein were considered. Thus, no significant differences in ciliation were found by either overexpression of ODF2/138NC and NEURL4, or recruitment of NEURL4 to the centrosome. However, we found a significant increase of ciliated cells in the total cell population to approximately 23% when FRB::ECFP::NEURL4 was expressed (ciliated cells/total cells counted: 135/509, 84/501, 122/501, three biological replicates. p=0.012588 *; Student’s T-test, two-sided, homoscedastic).

### Formation of higher order structures by interaction between HYLS1 and ODF2

Finally, we investigated the effect of HYLS1 on ciliogenesis. The coding sequence of *Hyls1* was cloned in frame into p*FRB::ECFP* giving plasmid p*FRB::ECFP::Hyls1*. Expression of FRB::ECFP::HYLS1 in NIH3T3 cells caused an increase of ciliated cells to approximately 19% (ciliated cells/total cells counted: 82/510, 97/515, 108/505, three biological replicates), compared to approximately 12% in untreated control cells (see above), which is statistically significant (Student’s T-test, two-tailed, homoscedastic. p=0.01356664*). Expression of both, 138NC::mRFP::FKBP and FRB::ECFP::HYLS1, caused ciliation in approximately 14.5% of transfected cells (ciliated transfected cells/total transfected cells counted: 11/104, 4/102, 30/103, three biological replicates), and in approximately 13% of transfected cells when dimerization was induced with rapamycin (ciliated transfected cells/total transfected cells counted: 11/107, 16/103, 11/102, three biological replicates). Thus, the recruitment of HYLS1 to the centriole by dimerization with ODF2 did not induce cilia formation. FRB::ECFP::HYLS1 showed cytoplasmic expression in NIH3T3 cells (Fig. 8a). Co-expression of both, 138NC::mRFP::FKBP and FRB::ECFP::HYLS1, indicated protein interaction. Strikingly, the formation of distinct ring-as well as tube-like structures were found that stained positive for both, the mRFP-autofluorescence of 138NC::mRFP::FKBP and the immune-decoration of HYLS1 mediated by incubation with anti-GFP antibodies (Fig. 8d-g, h-k, l-o). Tubes and rings had a diameter of approximately 560nm (calculated from 23 measurements taken from Fig. 8d and i). Higher-order structures were found in some of the transfected cells, while in some cells even colocalization to 138NC::mRFP::FKBP fibers could not be detected (Fig. 8p-s). The formation of tube-like structures and rings, which are most likely tubes in cross-section, was independent of rapamycin-induced dimerization suggesting that ODF2 and HYLS1 interact with each other to promote the formation of higher-order structures.

**Fig. 8.**
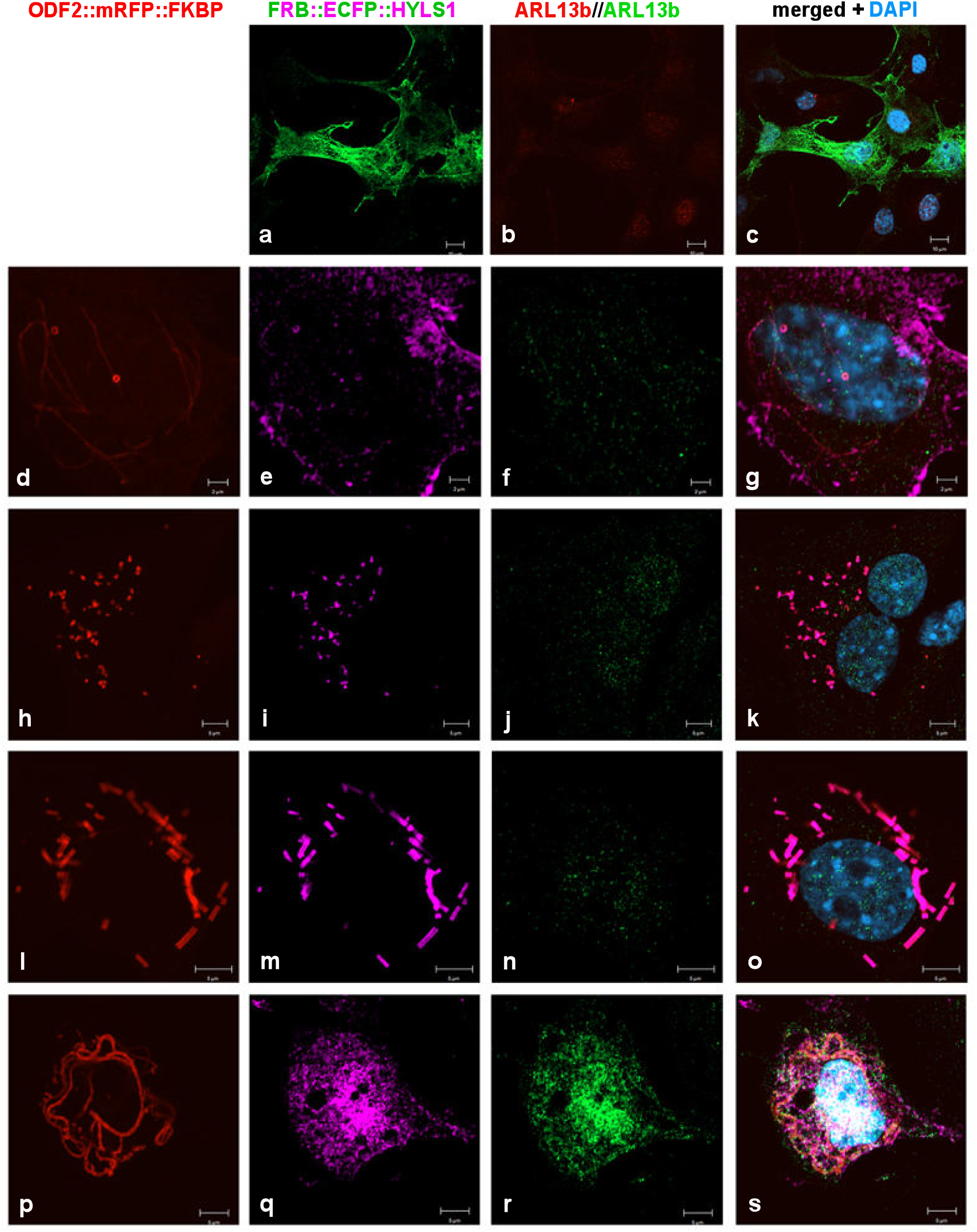
Formation of tubes and rings by interaction of ODF2 and HYLS1. (a-c) Cytoplasmic expression of FRB::ECFP::HYLS1. Anti-GFP immunostaining for the detection of FRB::ECFP::HYLS1 (green) and ARL13b (red). (d-g, h-k, l-o) Formation of higher-order structures, i.e. tubes and rings, by co-expression and co-localization of 138NC::mRFP::FKBP and FRB::ECFP::HYLS1. Autofluorescence of mRFP-tagged 138NC (red), and anti-GFP immunodecoration of FRB::ECFP::HYLS1 (pink) and ARL13b (green). (p-s) Formation of fibres by 138NC::mRFP::FKBP (red autofluorescence) without co-localization of FRB::ECFP::HYLS1 (pink). Anti-GFP immunostaining of FRB::ECFP::HYLS1 (pink) and ARL13b (green). Localization of 138NC::mRFP::FKBP (d, h, l, p), FRB::ECFP::HYLS1 (a, e, i, m, q), ARL13b (b, f, j, n,r), and merged images including nuclear stain with DAPI (blue) (c, g, k, o, s). Bars are 10µm (a-c), 2µm (d-g), and 5µm (h-s).

## Discussion

### Rapamycin-induced targeting to the centriole

The rapamycin-induced dimerization system has been widely used for the targeted recruitment of proteins (Putyrski and Schultz, 2012). We asked here, whether chemically induced dimerization can be used to control the location of centriolar proteins to modify the composition and function of centrioles. Specifically, we investigated whether the formation of primary cilia can be induced by the targeted recruitment of proteins that are essential for ciliary growth. We have chosen ODF2 as the binding partner for the recruitment of NEURL4 and HYLS1 because ODF2 is intrinsically located in the mother centriole and the basal body. NEURL4 and HYLS1 are essential for the elimination of the inhibitory cap, consisting of CP110, CEP97, CEP290, and other proteins, that is present at the distal ends of mature centrioles/basal bodies preventing axoneme extension (Spektor et al., 2007). We have fused the FK506 binding protein FKBP12 to ODF2 and the FRB-domain to either NEURL4 or HYLS1 and induced dimerization by rapamycin treatment. We show here as proof of principle the efficient recruitment of NEURL4 to the centriole by dimerization with ODF2 after mild rapamycin treatment.

### ODF2 controls CP110 levels

ODF2 is mandatory for the formation of primary cilia and facilitates the docking of ciliary vesicles to distal appendages (DAs) and the formation of the ciliary membrane via interaction with the GTPase membrane trafficking regulator Rab8a (Yoshimura et al., 2007; Wang et al., 2018; Sullenberger et al., 2020). Furthermore, the assembly of subdistal appendages (SDAs) depends on ODF2, which is located near the centriole wall. The formation of the primary cilium starts at the distal end of the mother centriole by the docking of ciliary vesicles to the tip of DAs and the concurrent extension of the ciliary axoneme (Sánchez and Dynlacht, 2016). DAs and SDAs are assembled by the sequential recruitment and positioning of specific proteins when the daughter centriole is converted into the mature or mother centriole (Yang et al., 2018; Chong et al., 2020). DAs are assembled by the orchestrated recruitment of CEP90, CCDC41/CEP83, CEP123/CEP89, SCLT1, CEP164, and FBF1, i.a., and assembly crucially depends on C2CD3, localized at the distal ends of centrioles (Tanos et al., 2013; Joo et al., 2013; Sillibourne et al., 2013; Wei et al., 2013; Ye et al., 2014; Kumar et al., 2021). CEP164 is positioned at the tip of DAs and initiates cilium formation by the docking of ciliary vesicles (Schmidt et al., 2012). However, an essential requirement for centriole maturation and appendage formation is the removal of DCPs (daughter centriole enriched proteins), a specific set of proteins including CEP120, Centrobin, and NEURL4 that are recruited to the nascent daughter centriole (Zou et al., 2005; Mahjoub et al., 2010; Li et al, 2012; Lin et al., 2013). DCP removal is regulated by TALPID3, located at the distal ends of both centrioles, and C2CD3, and TALPID3-deficiency abrogates DCP removal, assembly of DAs, and ciliogenesis, without affecting SDA assembly (Wang et al., 2018).

In addition, the inhibitory cap at the distal ends of centrioles consisting of CP110 and other proteins must be removed, which is achieved by degrading CP110 (Spektor et al., 2007; Tsang et al., 2008; Schmidt et al., 2009; Avidor-Reiss and Gopalakrishnan, 2013; Tsang and Dynlacht, 2013). CP110 is mainly degraded by the ubiquitin-dependent proteasome pathway in which the binding to the NEURL4-HERC2 complex plays an essential role (D’Angiolella et al., 2010; Al-Hakim et al., 2012; Li et al., 2013; Gonçalves et al., 2021). Targeting NEURL4 to the centrosome is sufficient to remove CP110 and restore ciliogenesis (Loukil et al., 2017). The transient translocation of NEURL4 to the mother centriole early during ciliogenesis requires the physical proximity between the mother and the daughter centrioles (Loukil et al., 2017). In summary, NEURL4 appears to play a role in both centriole maturation and axoneme extension. Removal of DCPs, including NEURL4, is the initial step in centriole maturation and appendage formation, while NEURL4 appears to be necessary for the degradation of the inhibitory cap at the distal end to allow the cilium to form. NEURL4, therefore, has to be recruited to the distal end of the mother centriole to promote CP110 degradation.

We have demonstrated the efficient recruitment of NEURL4 to the centriole by rapamycin-induced dimerization with ODF2. However, we found no indications that the targeted recruitment of NEURL4 to the distal end of the mature centriole promoted the loss of CP110. Although we found a significant decrease of CP110 at the basal body in ODF2 and NEURL4-transfected cells, the loss of CP110 was independent of rapamycin treatment. These results suggested that ODF2 may play an essential role in the elimination of CP110. To further substantiate, we depleted ODF2 by transfection of the short hairpin-plasmid *sh3* or *Odf2* siRNA and measured the amount of CP110 at the centrosome. We found an opposing trend in that a decrease in ODF2 correlated with an increase of CP110, and vice versa in the rescue experiments. Our data thus show that ODF2 is essential for the elimination of CP110.

The importance of NEURL4 for the degradation of CP110 is well documented (Loukil et al. 2017). Furthermore, the observation that the master regulator of the ubiquitin proteasome system, the deubiquitinase ubiquitin-specific protease-14 (UPS14), controls cilia formation and that generation of linear ubiquitin chains at CP110 by the linear ubiquitin chain assembly complex (LUBAC) is required for CP110 removal and ciliogenesis highlights the importance of ubiquitin-dependent proteolysis of CP110 (Massa et al., 2018; Shen et al., 2021). We found that ectopically expressed NEURL4 was localized to the centrioles without rapamycin treatment, suggesting intrinsic targeting of NEURL4 to the centrioles, but that rapamycin-mediated dimerization with ODF2 caused a large increase in the concentration of NEURL4 at the centrioles. Moreover, our data suggest that a specific amount of NEURL4 is sufficient to induce CP110 removal and that its further recruitment has no additional effect on CP110 degradation.

Although NEURL4 seems to play a major role in the ubiquitin-dependent proteolysis of CP110, it is unclear how NEURL4 is recruited, activated, or stabilized at the basal body prior to the extension of the ciliary axoneme. Our data indicated an involvement of ODF2 in this process.

We have shown that ODF2 plays an essential role in controlling the amount of CP110 at the centriole/basal body. This is further supported by the significant enrichment of ODF2 in basal bodies over centrosomes in both proliferating and serum-starved cells, and by the observation that a higher amount of ODF2 in the basal body correlated with a lower amount of CP110, and vice versa in the attached daughter centriole. ODF2 is a structural protein without any known enzymatic activity. Secondary structure predictions indicate that ODF2 is a protein composed mainly of alpha-helix structures, which can form coiled coils (Lupas et al., 1991; AlphaFold Monomer v2.0, Jumper et al., 2021; Varadi et al., 2021). Thus, ODF2, like many coiled-coil proteins, most likely acts in nucleating and scaffolding large macromolecular complexes (Truebestein and Leonard, 2016). More than 150 proteins interacting with ODF2 have been annotated in BioGRID4.4, including NEURL4 and CP110, both identified by proximity label-MS (Gupta 2015; BioGrid4.4., Stark et al., 2006). Our data show that the amounts of ODF2 and CP110 are interdependent, suggesting that ODF2 most likely acts as a scaffold for NEURL4 binding, which in turn promotes the degradation of CP110.

### ODF2 is not sufficient to promote cilia formation

We then asked whether overexpression of ODF2 might promote cilia formation via the removal of CP110. Either one of three different plasmids encoding ODF2 was transfected and the percentage of ciliated cells was determined. We found that the percentage of ciliated cells was in the same order of magnitude as in the non-transfected control cells. We, therefore, concluded that overexpression of ODF2 is not sufficient to promote cilia formation. We also found no significant increase in ciliated cells when ODF2 (*p138NC::mRFP::FKBP*) and NEURL4 (*FRB::ECFP::Neurl4*) were co-transfected and treated either without or with rapamycin. The percentage of ciliated cells was similar to that of the non-transfected control cells. However, overexpression of only NEURL4 resulted in a significant increase in ciliated cells, suggesting that increased levels of NEURL4 may increase the degradation of CP110. CP110 has a dual function as a suppressor and promoter of ciliogenesis, as it is also required for SDA assembly and cilia formation in vivo, suggesting that optimal cellular CP110 levels must be strictly balanced (Song et al., 2014; Yadav et al., 2016; Walentek et al., 2016). We, therefore, hypothesize that a moderate increase in NEURL4 would promote the degradation of CP110 to allow axoneme extension and cilium formation, whereas in the presence of excess ODF2, the amount of CP110 may have fallen below a certain threshold that prevents cilium formation.

Degradation of CP110 itself is not sufficient for cilium formation. In the G2-phase, in which only a few cells have a primary cilium, CP110 interacts with Cyclin F, which is the substrate-recognition subunit of the SCF ubiquitin ligase complex, on centrioles causing CP110 ubiquitylation and degradation by the proteasome (D’Angiolella et al., 2010). In addition, other CP110 binding partners are involved in the regulation of cilium assembly (Tsang and Dynlacht, 2013). The calcium-binding protein Centrin2, CETN2, an interactor of CP110, is essential for the efficient removal of CP110 and primary ciliogenesis upon serum starvation (Prosser and Morrison, 2015). Centrobin (CNTROB) is a component of the DCPs, enriched at the daughter centrioles, but also associates with the mother centriole upon serum deprivation and is necessary for primary ciliogenesis (Ogungbenro et al., 2018). Axoneme extension, in addition, requires the microtubule (MT)-associated protein/MT affinity regulating kinase 4 (MARK4) (Kuhns et al., 2013). MARK4 locates at the basal body and the ciliary axoneme. Depletion of MARK4 blocked ciliogenesis and prevented the exclusion of CP110/CEP97 from the mother centriole. MARK4 is activated by LKB1/STK11 recruiting ODF2 to the centriole and phosphorylating it. However, the effect of MARK4 on CP110 and cilia formation could be a purely indirect effect, as it primarily affects the recruitment of ODF2, which in turn promotes CP110 removal and cilia formation.

### Co-localization of ODF2 and HYLS1

The disappearance of CP110 from the basal body coincides with an enrichment of the serine/threonine kinase TTBK2, a microtubule plus-end tracking protein that phosphorylates not only Tau and Tubulin but also CEP164, CEP97, and MPP9 (Liao et al., 2015; Huang et al., 2018; Lo et al., 2019). TTBK2 is recruited to the centriole by CEP164 and that is required for the phosphorylation of CEP83, and the subsequent removal of CP110 and initiation of ciliogenesis (Goetz et al., 2012; Cajanek and Nigg, 2014; Oda et al., 2014; Lo et al., 2019). Binding of TTBK2 to CEP164 is regulated by phosphatidylinositol-4-phosphate (PtdIns(4)P) levels at the centrosome/ciliary base (Xu et al., 2016). Upon serum starvation, the phosphatidylinositol 5-phosphatase (INPP5E) departs from the centrosome causing depletion of PtdIns(4)P by the activity of phosphatidylinositol(4)P 5-kinase (PIPKIγ) thus freeing CEP164 for the binding with TTBK2 (Wang and Dynlacht, 2018). Furthermore, PIPKIγ recruits HYLS1 (hydrolethalus syndrome protein 1) to the ciliary base, which in turn causes activation of PIPKIγ, depletion of centrosomal PI(4)P, and axoneme extension (Chen et al., 2021). HYLS1 has been shown to play an essential role in cilia formation (Dammermann et al., 2009). Given the central role of HYLS1 in regulating PtdIns(4)P-levels at the ciliary base, recruitment of TTBK2, and removal of CP110, we asked here, whether chemically induced recruitment of HYLS1 to the centriole via dimerization with ODF2 promotes cilia formation. When ODF2 (*p138NC::mRFP::FKBP*) and HYLS1 (*FRB::ECFP::Hyls1*) were co-expressed in NIH3T3 cells, the percentage of ciliated cells was in the same order of magnitude as in the non-transfected control cells even when dimerization was induced by rapamycin-treatment. However, we observed a statistically significant increase in ciliation (p*) when HYLS1 was overexpressed, an observation also made for NEURL4-overexpression (see above). We, therefore, speculate that, similar to the case of NEURL4, a moderate increase in HYLS1 promotes cilia formation by increasing the removal of CP110, but that a further increase in HYLS1 at the centriole by targeted recruitment via ODF2 prevents cilia formation because CP110 is fallen below a certain threshold level. We observed co-localization of ODF2 and HYLS1 in the cytoplasm of transfected cells and their association into tubes and rings. Since tubes and rings were also observed in cells without rapamycin-induced dimerization, ODF2 most likely interacts with HYLS1 to promote the formation or stabilization of higher-order structures. The observed tubes and rings have an average diameter of approximately 560nm and are therefore most likely not centrioles, which have a diameter of only ∼250nm in vertebrate cells (Winey and O’Toole, 2014). The width of these structures is similar to the width of vertebrate sperm tails, which ranges from >640 to ∼1000nm, depending on the species and the section of the sperm tail (Menkveld et al., 2011; Weber et al., 2014). HYLS1 is highly expressed in testis (GeneCards) and is important for flagella formation in *Drosophila* spermatids (Hou et al., 2020). It, therefore, seems possible that the interaction between ODF2 and HYLS1 has promoted the formation of accessory structures similar to the outer dense fibers in the sperm tails in the cytoplasm of transfected cells and that their interaction may also be important for the formation of the outer dense fibers and the stability of the sperm tails.

### Conclusions

We used the rapamycin-induced dimerization system to efficiently recruit proteins to the centriole. We found that ODF2 is essential for the efficient removal of CP110. Since ODF2 is a structural protein implicated in the nucleation and scaffolding of macromolecular complexes, we hypothesize that ODF2 recruits and binds NEURL4 at the centriole, enabling the degradation of CP110. Forced expression of ODF2 with or without concomitant overexpression of its interacting proteins NEURL4 or HYLS1 did not stimulate cilia formation, whereas overexpression of either NEURL4 or HYLS1 resulted in a significant increase in ciliated cells, suggesting that CP110 levels must be tightly controlled to promote cilia formation.

## Materials and Methods

### Plasmid constructs

For targeted recruitment, the coding sequence of *Odf2,* subclone *138NC,* was ligated in frame to *mRFP-FKBP* at its 3’-end in the vector backbone of *pEGFP-N1*, which was deleted for the coding sequence of *egfp,* giving plasmid *p138NC(ODF2)::mRFP::FKBP*. 138NC targets the centrosome and lacks 173aa at its C-terminal end, otherwise present in human Cenexin (Q5BJF6.1) (Hüber et al., 2008). *Neurl4* was in frame ligated to FRB-ECFP at its N-terminal end in plasmid *pCR3.1* giving plasmid *pFRB::ECFP::Neurl4*. The full-length coding sequence of 962bp of *Hyls1* was isolated from mouse testis cDNA by RT-PCR using primer pair Hyls_NotI_rev (agcggccgcttaagaaggagaaagcgg)/ Hyls1_BglII_for (cagatctatggcagaaaaaagacaagc) and N-terminally ligated to FRB::ECFP in *pCR3.1* giving plasmid p*FRB::ECFP::Hyls1*. Centrin-2 fused to Cherry (*pCentrin-2::Cherry*) was transfected for identification of centrosomes by auto-fluorescence. Clones were sequenced to verify the correct open reading frame.

### Immune-cytology and protein quantification

NIH3T3 cells were obtained from DSMZ (ACC59) and cultivated in Dulbecco’s Modified Eagle’s Medium (DMEM; GlutaMax^TM^ with high glucose concentration (4.5g/l); #10566, ThermoFisher Scientific, Waltham, USA), supplemented with 1,000 U/ml penicillin and 1,000 µg/ml streptomycin and either 10% (v/v) fetal calf serum (FCS) (normal medium, NM) or 0.5% FCS (serum starvation medium, SSM). Cells were cultivated at 37°C and 5% CO_2_. Cultivation in serum starvation medium was for 24 hrs or 48 hrs.

For immune-cytology, cells were seeded at a density of 2 x 10^5^ cells per well of a 6-well plate on glass coverslips and fixed in 3.7% paraformaldehyde (PFA) for 20 min at 4°C. Cells were then permeabilized with 0.3% Triton X-100 in PBS (phosphate-buffered saline) for 10 min at room temperature and blocked by incubation in PBS containing 1% bovine serum albumin (BSA) and 0.5% Tween-20 for at least 1 hour. Samples were incubated with the primary antibodies anti-acetylated α-tubulin (clone 6-11B-1; Santa Cruz Biotechnology, Inc., #sc-23950, diluted 1:50), anti-ARL13B (#17711-1-AP, diluted 1:400, Proteintech, St. Leon-Rot, Germany), anti-ODF2 (ESAP15572, diluted 1:100, antibodies-online GmbH, Aachen, Germany), anti-CP110 (#12780-1-AP, diluted 1:100, Proteintech, St. Leon-Rot, Germany), or anti-GFP (either monoclonal mouse IgG2a (3E6), A-11120, lot #7101-1, Molecular Probes, Inc., Eugene, USA, or polyclonal rabbit anti-GFP, self-made) at 4°C overnight. Secondary antibodies used are goat anti-mouse-IgG-DyLight 488 (#35503, diluted 1:300, ThermoFisher Scientific, Waltham, USA), goat anti-rabbit-MFP590 (#MFP-A1037, diluted 1:100, Mobitec, Göttingen, Germany), goat anti-rabbit-IgG-Dylight 488 (#35553, ThermoFisher Scientific, Waltham, USA), and goat anti-rabbit-AlexaFluor-647 (#A-21245, lot AB_2535813, ThermoFisher Scientific, Waltham, USA) or goat anti-mouse-AlexaFluor-647 (#A-21235, lot AB_2535804, ThermoFisher Scientific, Waltham, USA). DNA was counterstained with DAPI (4’, 6-diamidino-2-phenylindole; Vector Lab., cat. no. H-1500). Images were taken by confocal microscopy (LSM 980, Carl Zeiss AG, Oberkochen, Germany) and processed using Adobe Photoshop 7.0. For protein quantification, all pictures were captured using identical settings, and intensities were measured using Fiji (Schindelin et al., 2012). The relative quantity of the protein in question was obtained by subtraction of the average intensity of the centrosomal/basal body background, using always four different areas for each quantification, from the intensity of the protein in question, followed by dividing the intensity through the area. Fold changes were calculated by relating the relative quantities to the average relative quantity of the control. The diameter of ODF2/HYLS1 structures was measured with Fiji. Counting of primary cilia was done by visual inspection and scanning through all focal planes.

### Transfection of cells

NIH3T3 cells were seeded at a density of 1.25×10^5^ on coverslips in 6-well plates. Transfection was done 24hrs post-seeding and cells were processed for immune-cytology 24 hrs later. Plasmid DNA or siRNA was transfected using EndoFectin^TM^ Max Transfection Reagent following the manufactureŕs instructions (#EF014, GeneCopoeia, Inc., Rockville, USA). For *Odf2* knockdown, the short hairpin constructs *sh3* (specifically targeting sequence gaactcctccaggagatac of mouse *Odf2/Cenexin*; Tylkowski et al., 2014) or *Odf2* siRNA (stealth siRNA ODF2MSS207236; final concentration 40nM; Life Technologies, Carlsbad, USA) were used. As controls, the plasmid *K07* (OriGene Technologies, Inc., Rockville, USA), which lacked homology with any known mRNA, or the control siRNA (siGenome Non-targeting siRNA #1; D-001210-01-05; target sequence UAGCGACUAAACACAUCAA; Dharmacon Lafayette, USA) were used. Additionally, for rescue, the expression plasmid encoding human Cenexin (*hCenexin*) (Soung et al., 2006) was co-transfected. Co-transfection of *pCentrin-2::Cherry* served for the identification of the centrosome in transfected cells.

### Rapamycin-induced dimerization

Cells were transfected with *p138NC::mRFP::FKBP* and either *pFRB::ECFP::Neurl4* or *pFRB::ECFP::Hyls1* and dimerization induced by incubation with rapamycin (#553210, 1µM stock solution in DMSO, Millipore, Burlington, USA). As a control, the cells were incubated with the same amount of DMSO, which was used as the solvent for rapamycin, for the indicated time. For quantification of CP110 at the basal bodies, cells were incubated with anti-CP110 antibodies (#12780-1-AP, diluted 1:100, Proteintech, St. Leon-Rot, Germany) followed by goat anti-rabbit-IgG-AlexaFluor-647 (#A-21245, lot AB_2535813, ThermoFisher Scientific, Waltham, USA). Basal bodies were identified by the decoration of the primary cilium with anti-acetylated tubulin antibodies followed by goat anti-mouse-IgG-Dylight 488 (#35503, ThermoFisher Scientific, Waltham, USA).

## Acknowledgments

Plasmid mRFP-FKBP and Lck-ECFP-FRB were kindly provided by Carsten Schultz (Heidelberg, Germany). Neurl4 was a kind gift of Bryan Dynlacht (New York, USA). Human Centrin-2 was from Jeff Salisbury (Rochester, USA).

## Data availability

Not applicable. There are no datasets generated during and/or analysed during the current study.

## Author’s contributions

Material preparation, data collection, and analyses were performed by Madeline Otto and Sigrid Hoyer-Fender. MO and SHF prepared the figures. SHF designed the experiments and wrote the manuscript. All authors commented on previous versions of the manuscript. All authors read and approved the final manuscript.

